# Operating principles of circular toggle polygons

**DOI:** 10.1101/2020.11.22.392951

**Authors:** Souvadra Hati, Atchuta Srinivas Duddu, Mohit Kumar Jolly

**Affiliations:** Centre for BioSystems Science and Engineering, Indian Institute of Science, Bangalore, India; Undergraduate Programme, Indian Institute of Science, Bangalore, India

**Keywords:** Toggle switch, multistability, cell-fate decision, design principles, coupled feedback loops

## Abstract

Decoding the dynamics of cellular decision-making and cell differentiation is a central question in cell and developmental biology. A common network motif involved in many cell-fate decisions is a mutually inhibitory feedback loop between two self-activating ‘master regulators’ A and B, also called as toggle switch. Typically, it can allow for three stable states – (high A, low B), (low A, high B) and (medium A, medium B). A toggle triad – three mutually repressing regulators A, B and C, i.e. three toggle switches arranged circularly (between A and B, between B and C, and between A and C) – can allow for six stable states: three ‘single positive’ and three ‘double positive’ ones. However, the operating principles of larger toggle polygons, i.e. toggle switches arranged circularly to form a polygon, remain unclear. Here, we simulate using both discrete and continuous methods the dynamics of different sized toggle polygons. We observed a pattern in their steady state frequency depending on whether the polygon was an even or odd numbered one. The even-numbered toggle polygons result in two dominant states with consecutive components of the network expressing alternating high and low levels. The odd-numbered toggle polygons, on the other hand, enable more number of states, usually twice the number of components with the states that follow ‘circular permutation’ patterns in their composition. Incorporating self-activations preserved these trends while increasing the frequency of multistability in the corresponding network. Our results offer insights into design principles of circular arrangement of regulatory units involved in cell-fate decision making, and can offer design strategies for synthesizing genetic circuits.

## Introduction

During embryonic development, cellular differentiation generates a diversity of cell types with varying characteristics and functions. Complex regulatory networks drive these cell-fate decisions; elucidating the design principles of these networks is a central theme in dynamical systems biology [1]. In the process of cellular decision-making, a pluripotent cell might exhibit more than one stable steady state (phenotype) in response to various external and internal factors, without any differences in genetic information (i.e. via differential expression of genes in different states or phenotypes). This feature is referred to as multi-stability (co-existence of multiple phenotypes) and it underlies the dynamics of regulatory networks involved in cell-fate decision-making during development [2,3]. Such multistable dynamics and consequent phenotypic changes has also been recently seen in disease progression [4,5] and in cellular reprogramming [6,7]. Thus, elucidating the dynamics of such multi-stable networks holds promise for advancing our understanding of embryonic development as well as the latest applications in synthetic biology [8,9] and regenerative therapies [10].

One of the most commonly observed network motif in cell-fate decisions is a ‘toggle switch’, i.e., two mutually repressing transcription factors A and B, each of which acts as a ‘master regulator’ for specific cell fate (**Fig 1A**) [11]. The mutual repression allows for the toggle switch to have two possible outcomes - (high A, low B) and (low A, high B), thus driving an ‘all-or-none’ response. Therefore, this network enables the progenitor cell to choose between two possible ‘sibling’ cell fates [12,13]. For instance, PU.1 and GATA1 form a toggle switch that drives hematopoietic stem cells to either a common myeloid progenitor (PU.1 high, GATA1 low) or an erythroid one (PU.1 low, GATA1 high) [14]. Also, in *E. coli*, the construction of a toggle switch exhibiting bistability and switching between the two states in response to external signals has driven an extensive design of synthetic genetic circuits [15]. In a toggle switch, it is common for the ‘master regulators’ to self-activate. When one or both master regulators of a toggle switch self-activate, it can enable a third stable state – a hybrid (medium A, medium B) state, often mapped on to the common progenitor state [1,16].

**Figure 1:**
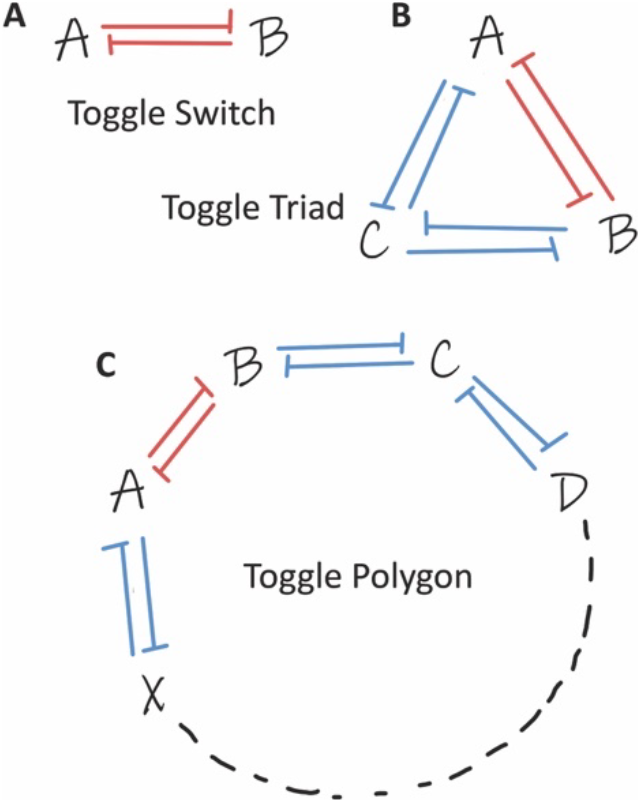
Schematics of toggle polygons. **A.** Toggle switch **B.** Toggle triad. **C.** A general toggle polygon with n nodes coupled circularly.

In cases when the same progenitor cells can give rise to more than two cell-fates, such as in T-cell differentiation, multiple toggle switches may be coupled. A regulatory network governing T-helper cell decisions to differentiate into Th1, Th2, and Th17 phenotypes includes a three-component network with master regulators of the three states (A, B and C) mutually repressing each other [17] (**Fig 1B**). Such a toggle triad, i.e., a circular coupling of three toggle switches (between A and B, between B and C, and between A and C), can enable three dominant distinct stable steady states - (high A, low B, low C), (low A, high B, low C) and (low A, low B, high C), each corresponding to a differentiated cell type. Similar to a toggle switch, including self-activations on A, B and C can enrich for hybrid states - (high A, high B, low C), (high A, low B, high C) and (low A, high B, high C) (Th1/Th2, Th2/Th17 and Th1/Th17 in this case) [18]. However, the dynamics and design principles of higher-order circular coupling of toggle switches remains unclear.

Here, we investigate the emergent dynamics of networks having the same functional units as toggle switch and toggle triad but with an increasing number of components. We have simulated, using both discrete and continuous simulations, different networks that are circular arrangements of toggle switches thus, we have named them ‘toggle polygons’ (**Fig 1C**). We noticed an intriguing pattern in their steady state distribution, depending on whether the toggle polygon is an even-numbered or an odd-numbered one. Even-numbered toggle polygons enable predominantly two states with consecutive components of the network expressing alternating high (1) and low (0) levels (1010…, 0101…). On the other hand, the odd-numbered toggle polygons enable more number of states – usually twice the number of components. Each of these states had comparable frequency and followed a ‘circular permutation’ pattern in their composition. Introduction of self-activations increased the multi-stability of the network without affecting the trends within the states enabled. Put together, our results unravel design principles of toggle polygons, i.e., networks including circular arrangement of toggle switches – the regulatory units involved in cell-fate decision making, and suggest strategies to design synthetic genetic circuits enabling such dynamic patterns.

## Results

### Even-numbered toggle polygons result in the two most frequent stable states

The dynamics of a toggle switch (**Fig 1A**) have been well-explored. It can give rise to two distinct phenotypes marked by expression levels (high A, low B) or (low A, high B), as witnessed in many scenarios during embryonic development. On a bifurcation diagram (or phase diagram), these phenotypes can exist independently (i.e., two monostable regions) or can co-exist (i.e., a bistable region) for a certain subset of parameter space [15,19]. Given the well-characterized dynamics of a toggle switch [20–23], including those that contain both microRNAs and transcription factors [24–28], we first investigated the dynamics of even-numbered toggle polygon (n = 4, 6, 8) (**Fig 2**).

**Figure 2:**
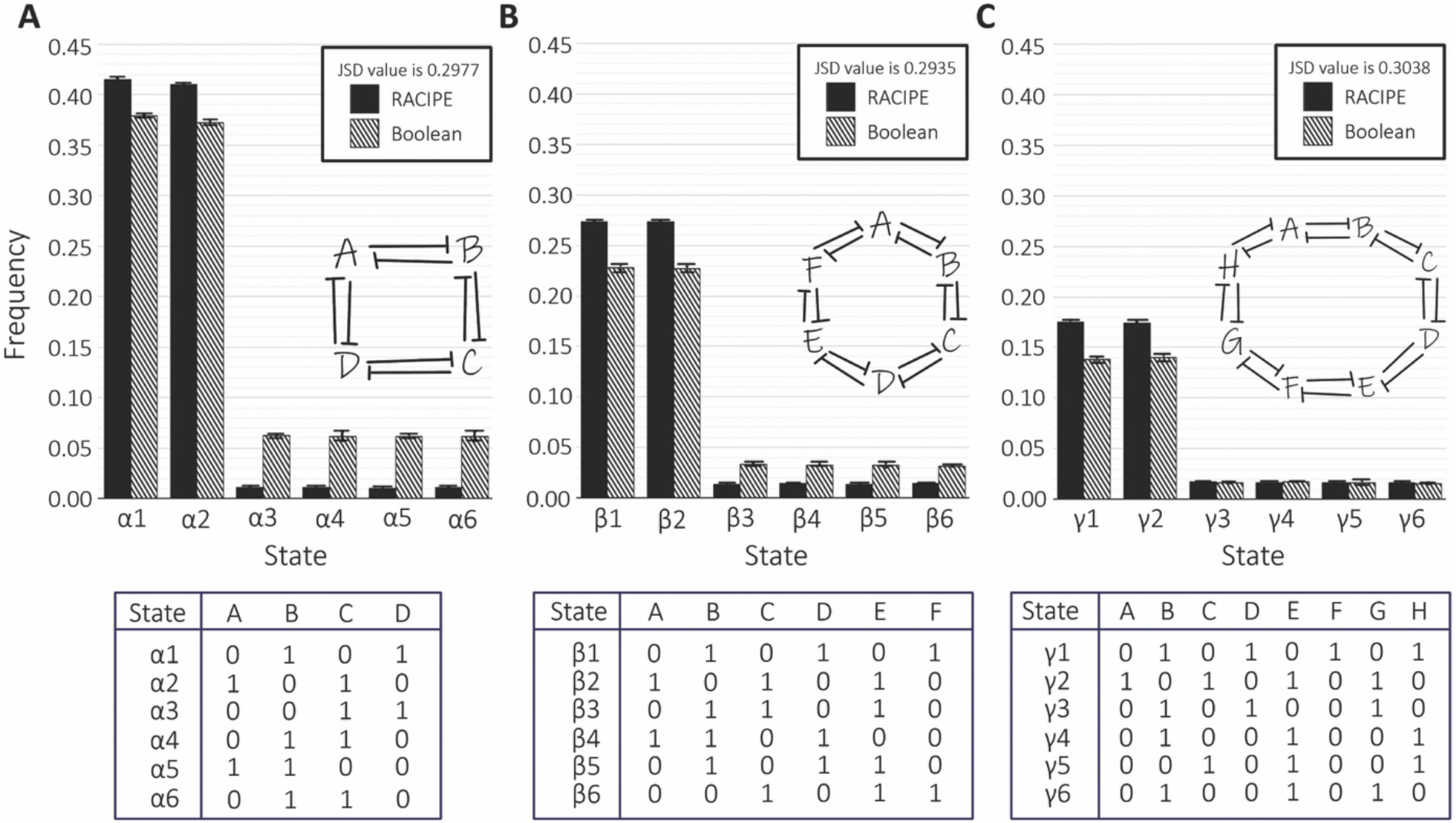
Even-numbered toggle polygons. Comparison between the most frequent solutions of the even-numbered toggle polygon networks as found from RACIPE and Boolean simulations. **A.** 4-component toggle polygon (toggle square) **B.** 6-component toggle polygon (toggle hexagon) **C.** 8-component toggle polygon (toggle octagon). For all the plots, n=3 independent RACIPE and Boolean replicates were done; error bars denote standard deviation. Labels 0 and 1 respectively represents lower and higher concentration of the corresponding component in the steady state solution of the corresponding state.

We simulated the dynamics of these networks using two complementary strategies – discrete and continuous – both of which take network topology as the input. For discrete simulations, we used a parameter-free approach: Boolean model using asynchronous update and Ising formalism [29]. For continuous simulations, we used a parameter-agnostic approach: RACIPE (Random Circuit Perturbation) that converts network topology information into a set of coupled ordinary differential equations (ODEs), samples 10000 parameter sets within a defined biologically relevant range for the given network topology, and identifies the steady states obtained for each parameter set for a varied set of initial conditions (see **Methods**) [30].

For the even-numbered toggle polygons (i.e., 4, 6, and 8 component networks), we observed two predominant stable steady states with equal frequencies, both via RACIPE and Boolean (**Fig 2 A-C**). These states had alternating high and low node expression levels. In a 4-component network, two states: (high A, low B, high C, low D) and (low A, high B, low C, high D) cumulatively have a frequency of 82% of the stable steady states obtained via RACIPE, and 75% in Boolean simulations (**Fig 2A**). This cumulative frequency shows a decreasing trend as the number of components in a toggle polygon increases. For the 6-component network, it is approximately 54% in RACIPE and 46% in Boolean simulations (**Fig 2B**). For the 8-component network, this frequency decreases to 35% and 28% for RACIPE and Boolean, respectively (**Fig 2C**).

Including self-activation to nodes in a network can affect the steady-state distribution. Thus, we ran RACIPE and Boolean simulations for the 4,6, and 8-component networks with self-activation (**Fig S1**). Similar to the case of toggle polygons without self-activation, these circuits showed a) two predominant states with both having the alternating high and low levels pattern and b) equal frequency of those two states. However, the cumulative frequency of two dominant states showed a slight decrease relative to the networks without self-activation (**Fig S1A-C**). Put together, these results suggest that a design principle of even-numbered toggle polygons is that they allow for two most predominant stable states, each of which shows alternating high and low levels of consecutive nodes in that toggle polygon. These trends are also congruent with pairwise correlations seen among different steady-state values of nodes in a network, as obtained via RACIPE simulations – while two consecutive nodes are negatively correlated, and alternative nodes are positively correlated. Usually, the longer the path between any two nodes, the weaker is the strength of correlation (**Fig S2,S3**).

To ensure that the results obtained for RACIPE are not due to under-sampling (i.e., the number of parameter sets sampled being insufficient), we performed RACIPE simulations for 25000 parameter sets, but did not observe any change in the qualitative features observed for steady-state distributions for ‘toggle hexagon’ (n=6) or ‘toggle octagon’ (n=8) with/without self-activation (**Fig S1D-G**). Finally, to quantify the similarity between the steady-state frequency distributions obtained from RACIPE and Boolean simulations for the circuits, we used an information-theoretic metric known as the Jensen-Shannon Divergence (JSD). JSD varies between 0 and 1; the higher the JSD value, the more dissimilar the two distributions are [31]. The relatively small values of JSD noted for even-numbered toggle polygons (**Fig1, S1**) reveal their robust dynamical features.

### Multistability in even-numbered toggle polygons

Next, we asked whether the abovementioned states can co-exist; in other words, can these toggle polygons be multistable for certain parameter sets. RACIPE simulates the dynamics of a given network for many initial conditions for a given parameter set. Thus, for every parameter set, we can identify whether it leads to monostable or multistable dynamics. Consistent with previous results [15,30], we noted that a toggle switch could be either monostable (~78%) or bistable (~22%) (**Fig 3A**). Moving from a toggle switch to a toggle square shows a drastic decrease in the frequency of parameter sets leading to monostability (~25%) and a consequent increase in those driving multistability, most evidently bistability (~65%). Consistently, as the number of components in a toggle polygon increased, the frequency of monostable parameter sets reduced, and that of multistable sets increased. This trend is consistent with our previous observations that the likelihood of multistability increases with an increase in the number of net positive feedback loops in a circuit [32] because higher even-numbered toggle polygons have a larger number of such feedback loops. There is one net positive feedback loop for a given node (say A) in a toggle switch (A inhibits B, which inhibits A). For node A in a toggle square, there are three positive feedback loops: two with adjacent nodes (A inhibits B, which inhibits A; and A inhibits D, which inhibits A), and one covering the entire circuit (A inhibits B, which inhibits C, which inhibits D, which inhibits A). Similarly, node A in the toggle hexagon and toggle octagon also has three positive feedback loops, thus increasing the chances of multistability compared to a toggle switch. This trend is reinforced by observations that incorporating self-activation can further decrease the frequency of monostable parameter sets, and increase that of multistable sets (**Fig 3B, S4A-C**).

**Figure 3:**
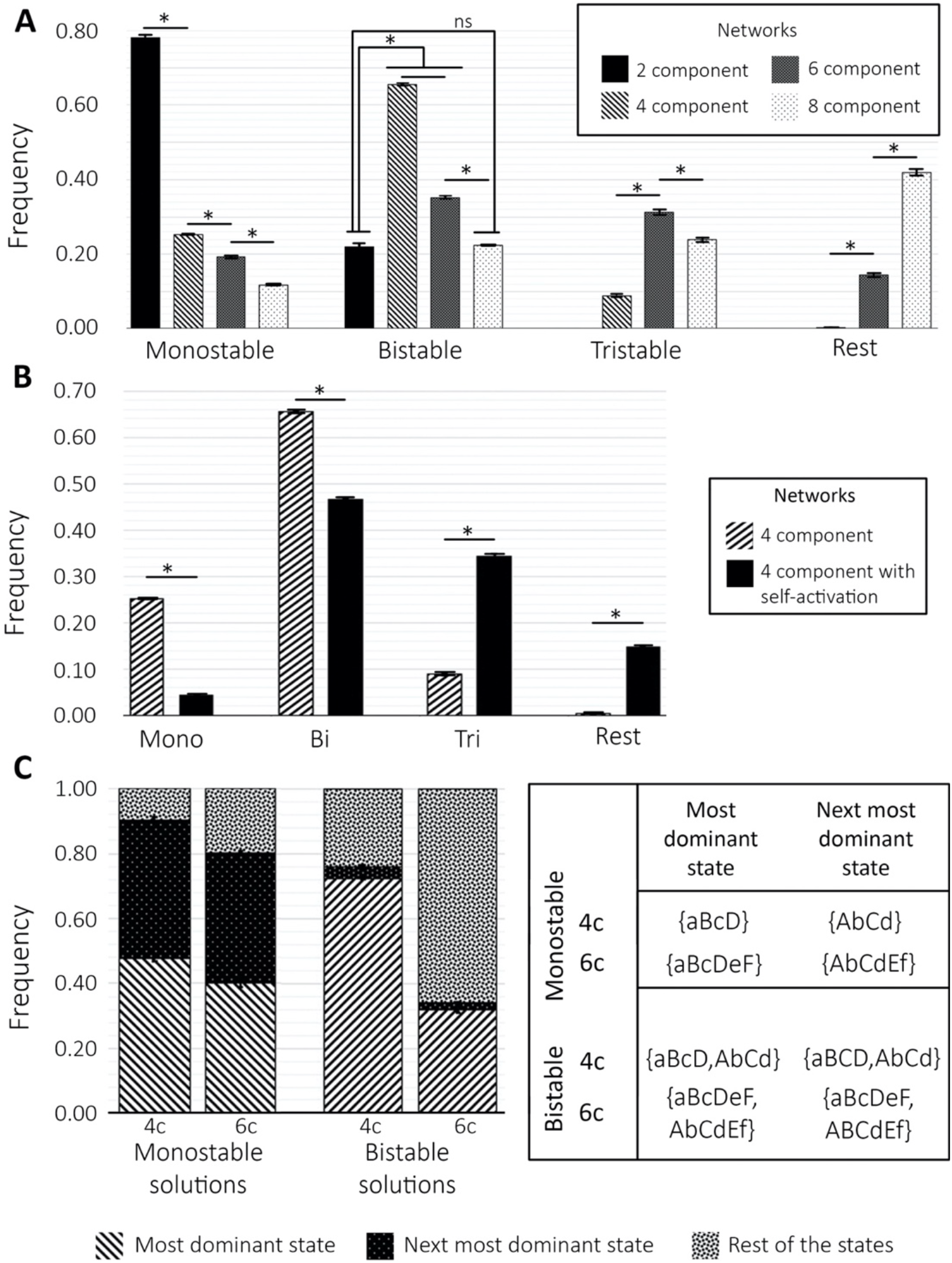
Multistability in even-numbered toggle polygons. **A.** Frequency of parameter sets in RACIPE that enable monostable, bistable and tristable solutions for different even-numbered toggle polygons - toggle switch (2 component/2c), toggle square (4 component/4c), toggle hexagon (6 component/6c) and toggle octagon (8 component/8c) networks. **B.** Comparison between frequencies of the monostable, bistable and tristable solutions form the RACIPE simulations of 4c and 4c with added self-activation on each node (4cS). **C.** Frequency of the most dominant state, the second most dominant state and rest of the states combined states (from bottom to the top respectively) in monostable and bistable solutions of RACIPE simulations for the 4c and 6c networks. For all the plots, n=3 independent RACIPE replicates were done; error bars denote standard deviation. * denotes p<0.01 for a Student’s t-test (‘ns’ implies p >0.01). Capital case letters indicate higher (1) level, small case letters indicate lower (0) levels of corresponding node in the corresponding steady state obtained. Thus, the notation AbCd means (high A, low B, high C, low D) or (A, B, C, D) = (1, 0, 1, 0).

Interestingly, as we move from the toggle square to the toggle hexagon, the frequencies of parameter sets for bistability are almost halved, but those corresponding to tristability tripled, and those enabling higher-order multistability (>=4 states) show an even stronger enrichment (**Fig 3A**). For a toggle octagon, the frequencies of the parameter sets driving bistability and tristability further decreased to about 23% each, while those driving higher orders of multistability increased to 42%. Overall, these results suggest that an increasing number of components in an even-numbered toggle polygon network can enable higher orders of multistability.

Further, we investigated the relative frequency of different possible states (or their combinations) in different monostable (or multistable) combinations. Among monostable states, the two most frequent states seen for toggle square were (high A, low B, high C, low D) ((A, B, C, D) = (1, 0, 1, 0) or {AbCd} henceforth) and (low A, high B, low C, high D) ((A, B, C, D) = (0, 1, 0, 1) or {aBcD} henceforth). Approximately 90% of parameter sets driving monostability led to either of these two states, with nearly 45% parameter sets corresponding to each (**Fig 3C**). Among bistable states, the phase containing a combination of these two states ({aBcD, AbCd}) was the most frequent (~75% parameter sets), suggesting that their co-existence was the most dominant form of bistability seen for a toggle square. Interestingly, the next dominant bistable set corresponding to only 2% of parameter sets. Similar results were observed for other even-numbered toggle polygons too. For instance, for a toggle hexagon – {aBcDeF} ((low A, high B, low C, high D, low E, high F)) and {AbCdEf} ((high A, low B, high C, low D, high E, low F)) – as the most frequent monostable states, and their co-existence formed the most frequent bistable case (**Fig 3C, S4D**).

Next, we investigated the relative stability of the two states that the network converges to for a bistable parameter set. For every parameter set that corresponded to the most dominant bistable phase corresponding to a network, we simulated 1000 initial conditions and counted how many initial conditions converged to state 1 (say x) and how many converged to state 2 (= 1000 - x). We plotted the distributions of x and (1000-x) taken over all parameter sets corresponding to this bistable phase. For a 4-component toggle polygon network (a toggle square), the most dominant bistable phase is ({AbCd, aBcD}). The distribution of the values of x (and that of 1000-x as well) drawn reveal two dominant extreme regions where x > 0.955 or x < 0.045 (i.e. 1000 – x > 0.955). For a toggle square, a significant part (~48%) of parameter sets belonged to these extreme regions. Thus, these parameter sets showed a stark difference in the relative stability of basins of attraction corresponding to the two states (**Fig 4A, 4D**). This observation suggested that a percentage of bistable parameter sets were effectively monostable because more than 95.5% of the sampled initial conditions led to only one of the two states. This trend was consistent for toggle hexagon with/without self-activation (**Fig 4B, 4D**). Incorporating self-activation pushed the distribution even further to the extremes for both the toggle square (frequency in the extreme regions = 64% for the case with self-activation) and the toggle hexagon (frequency in the extreme regions = 80% for the case with self-activation). Moreover, the larger the toggle polygon, the higher the frequency of such extreme parameter sets. Thus, while consistent trends were seen for toggle switch, for a toggle switch without self-activation, the distribution was not as extreme as seen for other networks (**Fig 4C**). Therefore, we concluded that while even-numbered toggle polygons did allow for multistability, many parameter sets leading to bistability can be asymmetric in terms of the relative stability of the two most dominant states seen.

**Figure 4:**
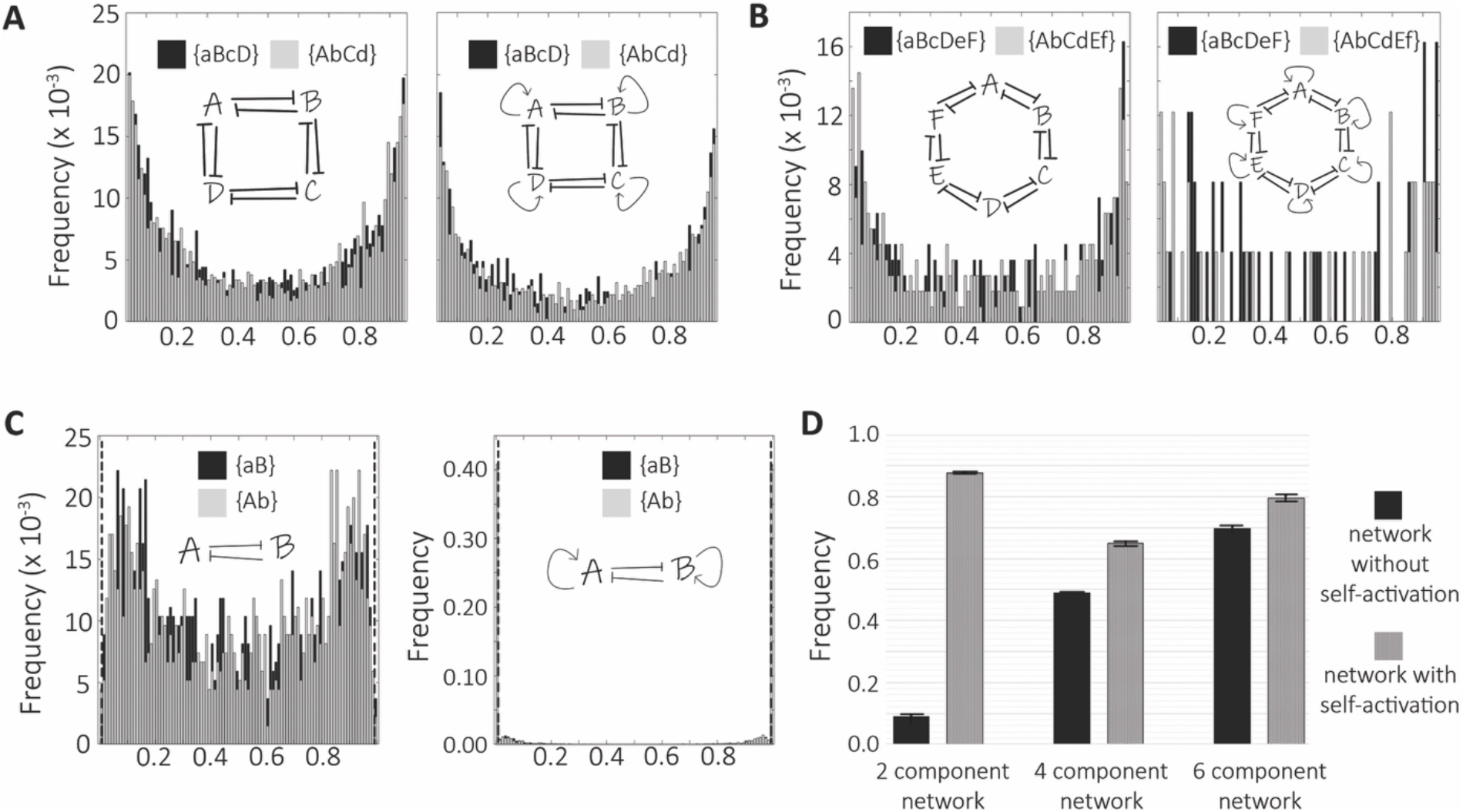
Relative stability of states in the most dominant bistable region. **A.** Distribution of the frequency of achieving the two states in the most dominant bistable solutions for RACIPE generated parameter sets for a toggle square with/without self-activation (4c, 4cS) (x-axis: 0.045-0.0955; frequency = no. of initial conditions converging to that state /total no. of initial conditions simulated for that parameter set (1000 in this case). **B.** Same as A. but for 6c and 6cS **C.** Same as A but for 2c and 2cS (x-axis: 0.00–1.00). Dotted vertical lines at the both the ends of the plots represents the boundary of 0.045 and 0.0955 considered as the region of effective monostable solutions. **D.** Probability of the frequency of the most dominant solution parameters belonging to “effective monostable” state (i.e. [0.00-0.455) U (0.095-1.00]) in each of the beforementioned networks. For all plots, n = 3 independent parameter sets were chosen from RACIPE, simulations of relative stability were done using the ode45 function in MATLAB (MathWorks Inc.); errors bars denote standard deviation.

### Design principles of odd-numbered toggle polygons

Next, we focused on odd-numbered toggle polygons. Our previous simulations showed that a toggle triad network could converge to one of the six dominant states: three ‘single-positive’ ones (only one of the nodes of the network is high, other two are low – (A, B, C) = (1, 0, 0), (0, 1, 0) or (0, 0, 1)) and the three ‘double positive’ ones (two of the nodes in the network are high – (A, B, C) = (1, 0, 1), (0, 1, 1), (1, 0, 1)) [18]. We next analyzed the steady states of higher order odd-numbered toggle polygon networks to find similarities and differences with trends seen for toggle triad. The 5-component toggle polygon network (toggle pentagon) is seen to have ten dominant states (**Fig 5A**), divided into two groups similar to the ‘single positive’ and ‘double positive’ states of the toggle triad. For a toggle pentagon, the dominant states of the two groups could be termed ‘double positive’ (two nodes in the network are high) and ‘triple positive’ (three nodes in the network are high) states. The five ‘double positive’ states have a frequency of about 10.5% each, while the ‘triple positive’ states have a frequency of about 8% according to RACIPE simulations, consistent with results for toggle triad that ‘single positive’ had higher frequency than the ‘double positive’ ones. On the other hand, Boolean simulation does not distinguish between these two groups, and all ten dominant states have a frequency of about 10% each. Consistent results were observed for toggle pentagon (7-component network) – 14 states in total and the seven ‘triple positive’ states were slightly more frequent than the ‘quadra positive’ states, based on RACIPE simulations (**Fig 5B**).

**Figure 5:**
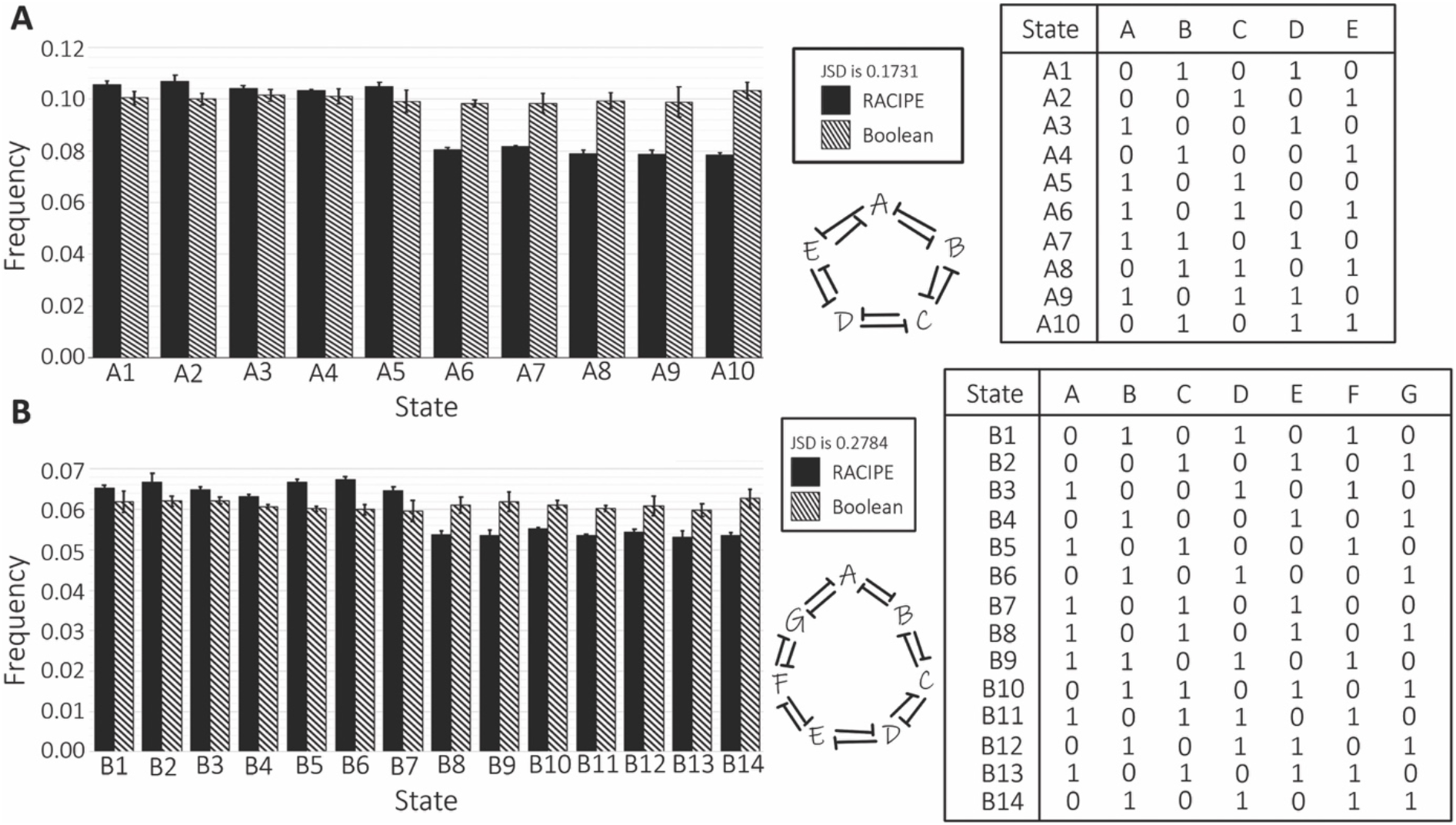
Odd-numbered toggle polygons. Comparison between the most frequent solutions of the different odd-element toggle polygon networks seen in RACIPE and Boolean simulations. **A.** 5 element toggle polygon (5c; toggle pentagon). **B.** 7 element toggle polygon (7c; toggle heptagon). 0 and 1 respectively represents lower and higher concentration of the corresponding component in the steady state solution of the corresponding state. For all plots, n=3 independent RACIPE or Boolean replicates were done; error bars denote standard deviation. JSD denotes Jensen-Shannon Divergence values as reported earlier.

Further, we investigated the influence of including self-activation. No significant qualitative changes were observed for toggle pentagon, toggle heptagon, consistent with our observations for even-numbered polygons (**Fig S5**). Consistently, toggle nonagon (9-component network) with/without self-activations led to 18 most frequent states – nine of which are ‘quadra positive’, and the remaining ones are ‘penta positive’, with the former ones slightly more abundance than the latter (**Fig S6**). Put together, a design principle of ‘n’-component odd-numbered toggle polygon is that it leads to ‘2n’ dominant states that follow the alternative high and low values of consecutive nodes as much as possible. Because of odd number of nodes, the alternate high/low pattern cannot be followed as coherently as for even-numbered polygon; thus, for states seen in odd-numbered polygons, two consecutive nodes tend to have the same value – 1 (high) or 0 (low).

Another feature observed for odd-numbered toggle polygons is that the ‘2n’ states are divided into two groups of ‘n’ each, with one group of states having slightly higher frequency relative to the other. The more prevalent states typically have fewer nodes being at a high value (or 1) as compared to the other set. A potential reason underlying this trend may be that the stable states with two adjacent node values being high (1) may be less frustrated [33] than those with two adjacent nodes being low (0).

Next, we investigated multistability enabled by odd-numbered toggle polygons. Reminiscent of our observations for even-numbered polygons, the frequency of parameter sets leading to monostability decreased, and consequently, those leading to multistability increased, as we studied higher-order polygons (from toggle triad to toggle pentagon to toggle heptagon) (**Fig 6A**). For the 5-component network, most parameter sets are bistable, but for the 7 and 9-component networks, there is a characteristic shift to higher-order multistability as seen for even-numbered polygons too. Apart from the increase in nodes, another trait that leads to a shift towards higher-order multistability is the incorporation of self-activation (**Fig S7**). For the 5, 7, and 9-component networks, the frequency of states of the order of multistability greater than or equal to three is 35%, 55%, and 70%, respectively.

**Figure 6:**
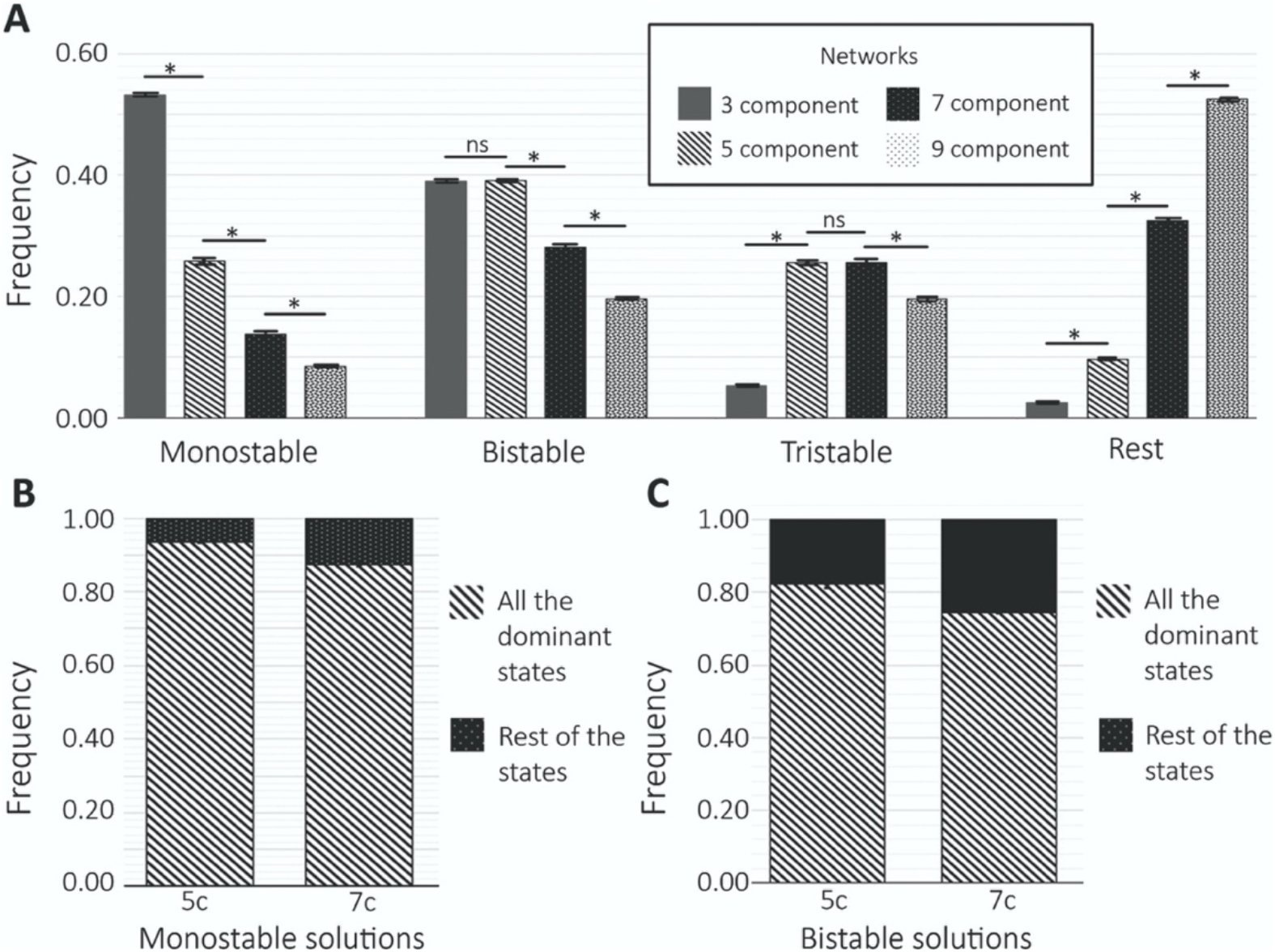
Multistability in odd-numbered toggle polygon networks. **A.** Frequency of parameter sets generated by RACIPE that enable monostable, bistable and tristable solutions for toggle triad (3c), 5c, 7c and 9c networks. **B.** Frequency of all the dominant monostable states combined with respect to the whole monostable solution space for the RACIPE simulations of 5c and 7c networks. **C.** Combined frequency of all the bistable states that are combinations of the most dominant monostable states with respect to the whole bistable state solution space for RACIPE simulations of 5c and 7c. n= 3 independent RACIPE replicates were done; error bars denote standard deviation. * denotes p<0.01 for a Student’s t-test (‘ns’ implies p >0.01).

Further, we examined the distribution of all monostable states driven by corresponding parameter sets. For the 5-component toggle polygon, the ten dominant states account for more than 90% monostable solutions. This dominance is seen for 7-component (**Fig 6B**) and other odd-numbered networks too (**Fig S8A**). Similarly, distribution within the bistable states is broadly conserved: for the 5-component network, all bistable phases (the combination of steady states) formed as a combination of any of the ten dominant monostable states together have a frequency of about 80% (**Fig 6B**). The dominance of these bistable states as a combination of any of the dominant monostable states continues for 7-component (**Fig 6B**) and, 9-component (**Fig S6A**) toggle polygon as well, thus resonating with our previous observation for the toggle triad network [18].

A closer analysis of the dominant steady-state solutions seen for odd-numbered toggle polygons reveals intriguing patterns. Consider the case of toggle pentagon. First, the ten states constitute five pairs of states; each pair has two states that are ‘mirror images’ of one another, i.e., one state can be obtained from another one by replacing every 0 with 1 and vice-versa in terms of node values; for instance, (A, B, C, D, E) = (1, 0, 1, 0, 1) and (0, 1, 0, 1, 0) are mirror images. This trend explains why we see five ‘double positive’ and five ‘triple positive’ states for a toggle pentagon. Second, if we consider all states, they are ‘circularly permutated’, i.e., one state can be obtained from another by taking the last node’s value in the sequence and appending it as the first node. Thus, from (1, 0, 1, 0, 1), if we move 1 to the beginning (and eliminate the consequent last node value), we get (1, 1, 0, 1, 0), which is also one of the steady-state solutions obtained. This state can be further ‘circularly permutated’ to give (0, 1, 1, 0, 1), following which we can get (1, 0, 1, 1, 0) and consequently (0, 1, 0, 1, 1) which further leads to (1, 0, 1, 0, 1), i.e. the initial state we started with, thereby completing the ‘circular permutation’.

For a toggle polygon, we also performed relatively stability analysis for few bistable phases, and observed similar pattern as seen for bistable parameter sets for even-numbered toggle polygons (**Fig S9**). Similarly, the correlation matrices of node values for results obtained through RACIPE reveal similar trends – strong negative correlation with consecutive nodes (**Fig S10, S11**) for a toggle pentagon and other odd-numbered toggle polygons with/without self-activation. These trends reveal some common trends in solutions obtained for both odd and even toggle polygons.

## Discussion

Cellular decision-making involves various network motifs, each with a specific set of functions [34]. One such ubiquitous motif is a toggle switch, i.e., a mutually inhibitory feedback loop between two components [1]. Such feedback loops can also be observed in cell-cell communication. One canonical example is Notch-Delta signaling [35–37] that can drive tissue patterning outcomes across biological contexts [38]. The dynamics of a toggle switch can be influenced by gene expression noise [39], epigenetic changes [40], partitioning errors during cell division [41–44], and conformational noise in intrinsically disordered regions of a component in a switch [45]. Thus, the deterministic and stochastic dynamics of toggle switches and coupling of many positive and negative feedback loops have been well-explored in many biological systems [19,20,46–51], including spatially extended realizations such as diffusion of mutually inhibiting molecules [52], and synthetic design of circuits to achieve a set of desired functions [53,54].

Here, we investigate the dynamics of toggle switches coupled circularly to form toggle polygons. We observed that even-numbered toggle polygons enable two most common states – the states where the expression of two consecutive nodes is anti-correlated among themselves (101010…., 01010…) (“1” indicates a relatively higher level, “0” indicates a relatively lower level). These results suggest that nodes in an even-numbered toggle polygon can follow the ‘salt-and-pepper’ pattern, reminiscent of patterns seen in a sheet of cells communicating via Notch-Delta signaling [55]. These patterns were conserved upon incorporating self-activation on the nodes in a toggle polygon. Therefore, the agreement in frequency distributions obtained for the even-numbered toggle polygons using both approaches – logical/Boolean models and continuous (RACIPE) – suggests this pattern as a design principle of such networks and offer a scaffold for synthetic network design enabling such outputs (**Fig 7**).

**Figure 7:**
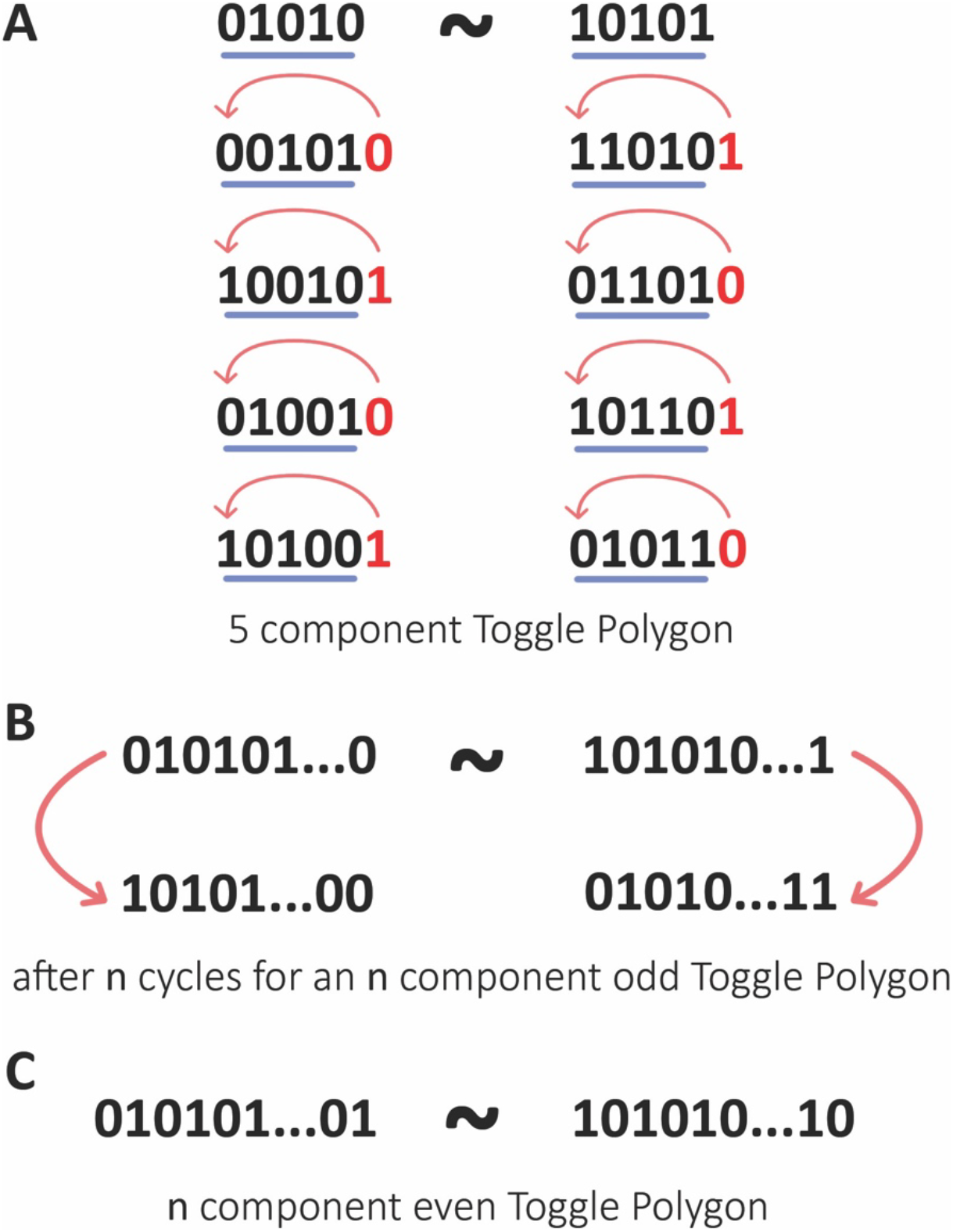
Patterns seen in the steady-state solutions obtained from toggle polygons. **A.** Schematic representation of the “complemented circular permutation” pattern shown by the most dominant monostable states of toggle polygon network. **A.** 5c network. **B.** General odd element toggle polygon network. **C.** General even element toggle polygon network, showing two most dominant states, each of which is a ‘mirror image’ of another one, and shows an alternative ‘salt-and-pepper’ pattern in terms of consecutive nodes being high (1) or low (0).

In contrast to even-numbered toggle polygons, odd-numbered toggle polygons show a much larger number of states, typically twice the number of elements in a toggle polygon. Nonetheless, these states also exhibit the ‘salt-and-pepper’ pattern (10101…, 01010…) as much as possible. For instance, a toggle triad can enable six states – three ‘single positive’ states (100, 010, 001) and three ‘double positive’ ones, which are ‘mirror images’ of the three abovementioned states (011, 101, 110). Similarly, a toggle pentagon may enable ten states – five ‘double positive’ states (10100, 01010, 00101, 10010, 01001) and five ‘triple positive’ ones that are ‘mirror images’ of the ‘double positive’ ones (01011, 10101, 11010, 01101, 10110) (**Fig 7**). The states in odd-numbered polygons appear more ‘frustrated’ than those seen in the even-numbered ones [33], and one set of states in odd-numbered polygons looks more ‘frustrated’ than the other one.

While toggle polygons for n > 3 are not necessarily explicitly identified yet in biological scenarios, a bidirectional coupling of two toggle switches similar to that of a toggle square has been observed in scenarios showing interconnected decision-making along two different axes of cellular plasticity [56]. The steady states obtained in this case are resonant with those reported here for a toggle square. Put together, our study demonstrates operating principles of toggle polygons, and reveal that odd-numbered and even-numbered polygons behave differently. These dynamical patterns seen suggest strategies for designing synthetic genetic circuits to generate this set of states.

## Author contributions

SH performed research and analyzed data, ASD analyzed data and wrote the initial draft of the manuscript, MKJ designed and supervised research and edited the draft.

## Acknowledgements

This work was supported by Ramanujan Fellowship awarded to MKJ by Science and Engineering (SERB), Department of Science and Technology, Government of India (SB/S2/RJN-049/2018).

## Materials and Methods

### RACIPE (<u>*Ra*</u>ndom <u>*CI*</u>rcuit <u>*Pe*</u>rturbation analysis)

RACIPE is a computational tool for investigating the dynamics of a given transcriptional network topology. It takes as input a network topology and converts it into a set of ordinary differential equations (ODEs) representing the interactions in that topology. For this set of ODEs, it samples a given number of kinetic parameters (10000 in this case) from a biologically relevant range. For each parameter set, multiple initial conditions (1000 in this case) are chosen randomly and the system is simulated to identify steady-state values for each component of the network. For a given parameter set, the system is classified as mono-, bi-, tri-stable to deca-stable depending on the number of steady states the set of initial conditions converge to. A generic ODE denoting the effect of component B on component A, as defined in the network topology file, denoted by RACIPE is:

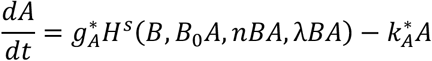

where, *g_A_*: production rate, k_A_: degradation rate, *H^S^*(*B*, *B*_0_*A*, *nBA*, *λBA*): shifted Hill function[1]

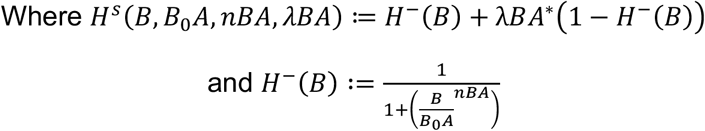

The steady solution generated by RACIPE is in log2 scale. For our analysis, we have normalized the steady state solutions obtained in two steps. First, we have performed g/k normalization for accounting for the extremes in sampling of the production and degradation rate parameters. g/k normalization includes dividing every steady-state value (E_i_) in the solution file by the ratio of the production and degradation rate of the respective component (g_i_/k_i_) of the network of the corresponding parameter set. Secondly, we have performed z-score normalization by calculating the mean and the standard deviation for every component ‘i’ over all parameter sets after the g/k normalization. The final transformation formula for each iteration looks like:

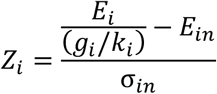

Here, *Z_i_*: final normalized expression, *E_in_*: mean, *σ_in_*: standard deviation

We have defined the states as ‘high’ and ‘low’ with respect to the normalized values being greater than and smaller than 0 respectively. For every network shown in the main text and supplementary material, three independent replicates of RACIPE simulation were performed and the analyzed data has been presented in the form of mean ± standard deviation.

### Boolean Analysis

For Boolean analysis, network topology is given as the input file that mentions the nodes and edges of the network. The edges can be of two types, activating and inhibiting. The analysis was carried out by the asynchronous update of the nodes, i.e. one node is chosen randomly and updated in a given timestep. The constraint of equal weightage to inhibitory and activating links was used. The updating of the nodes follows a simple majority rule (also called as Ising model). The node is updated to 1 if the sum of activations to the node is higher than inhibitions and updated to 0 for the opposite case. The steady state is said to be reached if there is no change in the updates for a predefined number of time-steps. We have run the simulations for 10000 random initial conditions for a given network.

### Jensen-Shannon divergence

The Jensen-Shannon divergence (JSD) values were calculated using the ‘Jensen Shannon’ function of SciPy module of Python [57].

### Relative Stability Analysis

The .prs file generated by RACIPE has the details of the parameter for each solution of RACIPE simulation. We identified parameter sets leading to more than one solution, wrote a typical ODE as that in RACIPE framework using these parameter sets, and for each parameter set, we counted how many initial conditions out of the 1000 randomly chosen ones converged to which of the reported states, to quantify the relative abundance of the two state (we looked at bistable cases). This process was repeated for many bistable parameter sets.

