## Supplementary Information for "Operating principles of circular toggle polygons"

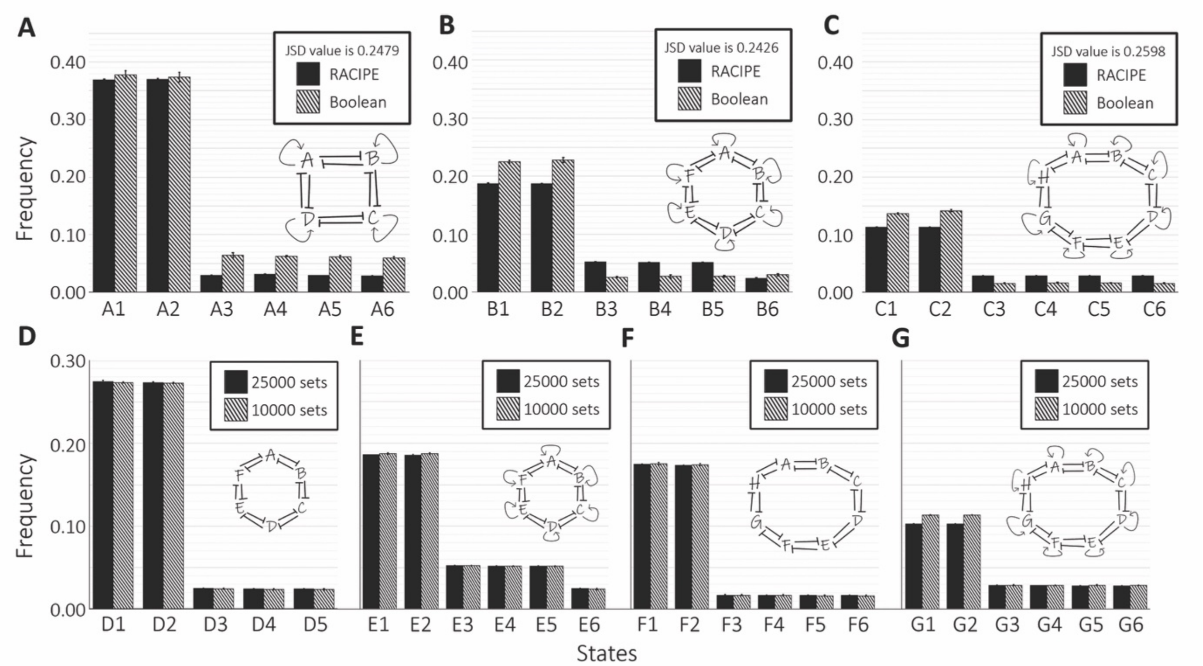


**Figure S1**: **Even-numbered polygons**. Comparison of the most frequent solutions of even element toggle polygon networks with added self-activation on each node as found from the RACIPE and Boolean simulations. **A.** 4c with self-activation *(4cS)*. **B.** 6c with self-activation *(6cS).* **C.** 8c with self-activation *(8cS)*. Comparison between the most frequent solutions of the large even element toggle polygon networks as represented by the RACIPE simulations with 1000 initial conditions and 25000 and 10000 parameter sets, respectively. **D.** 6c network. **E.** 6cS network. **F.** 8c network. **G.** 8cS network. We performed n=3 independent RACIPE simulations; error bars represent standard deviation.

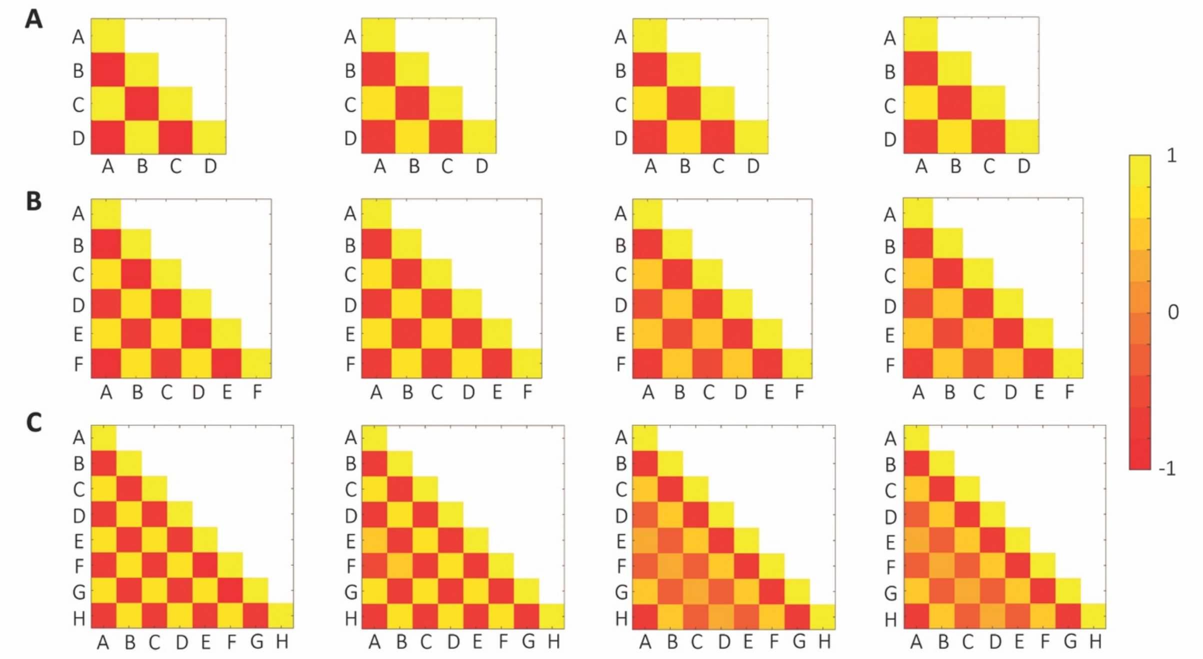


**Figure S2: Pairwise correlation matrices for even-numbered toggle polygons.** (Left to right) Pearson correlation coefficients of monostable RACIPE solutions, Spearman correlation coefficients of monostable RACIPE solutions, Pearson correlation coefficients of all RACIPE solutions combined, and Spearman correlation coefficient of all RACIPE solutions combined on circuit topologies **A.** 4c, **B.** 6c, **C.** 8cS. Each cell in this triangular matrix represents pairwise correlation. Colorbar shown to the left shows the value of correlation coefficients.

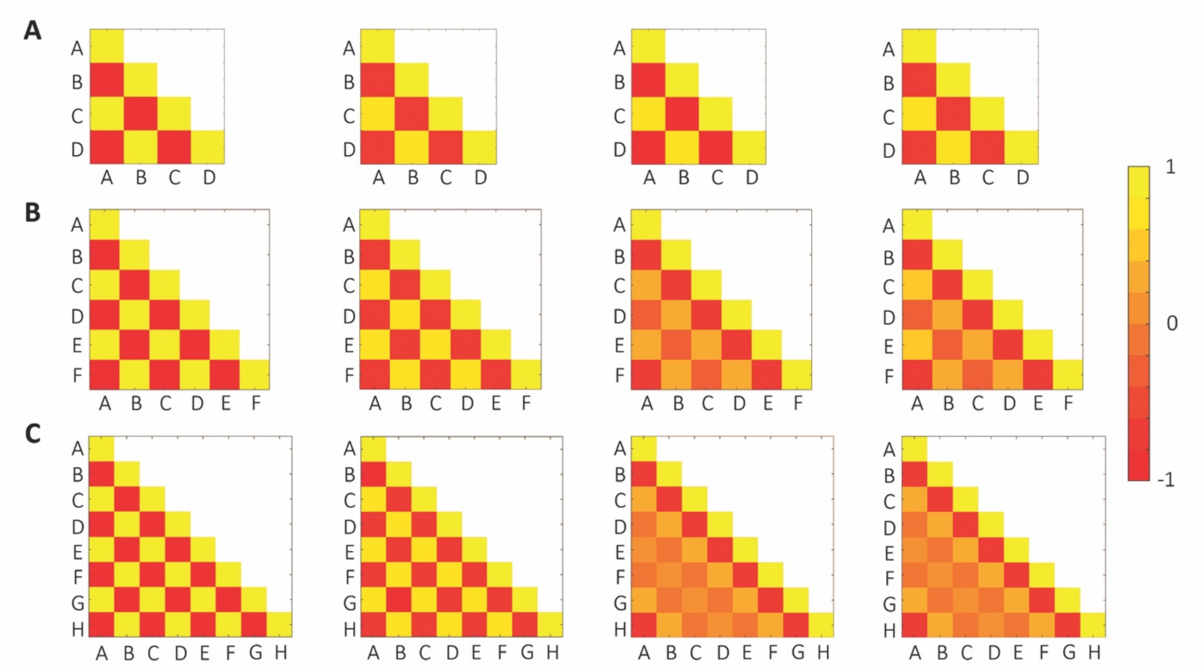


**Figure S3: Pairwise correlation matrices for even-numbered toggle polygons with self-activation.** Same as Fig S2 but for **A.** 4cS (toggle square with self-activation), **B.** 6cS (toggle hexagon with self-activation), **C.** 8c (toggle octagon with self-activation).

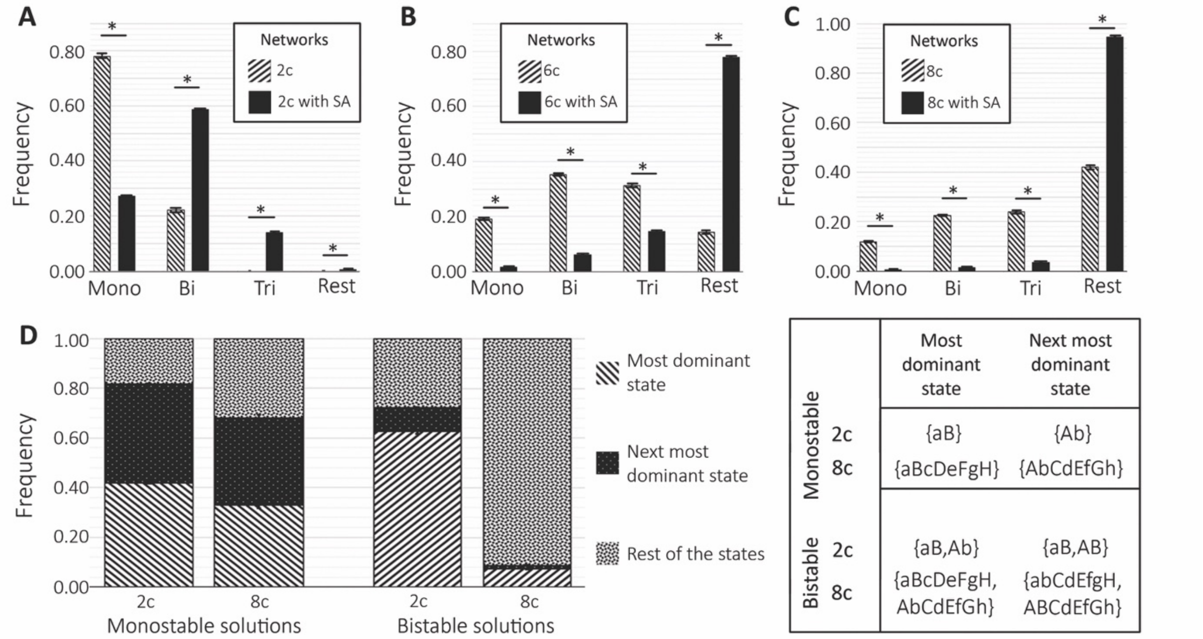


**Figure S4: Multistability in even-numbered toggle polygons**. Comparison between frequencies of monostable, bistable and tristable solutions form the RACIPE simulations of the even-numbered element toggle polygon networks with and without added self-activation at each node. **A.** 2c and 2cS networks. **B.** 6c and 6cS networks. **C.** 8c and 8cS networks. **D.** Frequency of the most dominant state, next most dominant state and rest of the combined states (from bottom to the top respectively) in monostable and bistable solutions of the RACIPE simulations of 2c and 8c networks. N=3 independent RACIPE replicated were done; error bars denote standard deviation. * $p<0.01$ for a Student's t-test, ('ns' implies $p > 0.01$).

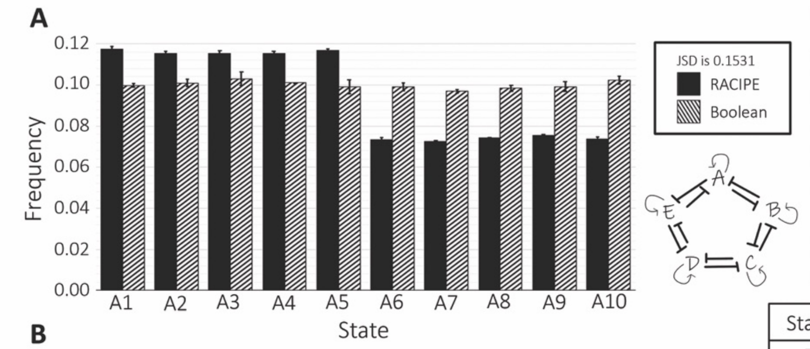


| State | A | B | C | D | E |
|---|---|---|---|---|---|
| A1 | 0 | 1 | 0 | 1 | 0 |
| A2 | 0 | 0 | 1 | 0 | 1 |
| A3 | 1 | 0 | 0 | 1 | 0 |
| A4 | 0 | 1 | 0 | 0 | 1 |
| A5 | 1 | 0 | 1 | 0 | 0 |
| A6 | 1 | 0 | 1 | 0 | 1 |
| A7 | 1 | 1 | 0 | 1 | 0 |
| A8 | 0 | 1 | 1 | 0 | 1 |
| A9 | 1 | 0 | 1 | 1 | 0 |
| A10 | 0 | 1 | 0 | 1 | 1 |

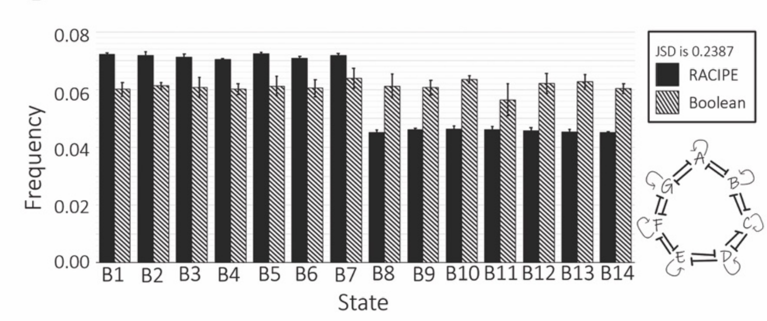


| State | A | B | C | D | E | F | G |
|---|---|---|---|---|---|---|---|
| B1 | 0 | 1 | 0 | 1 | 0 | 1 | 0 |
| B2 | 0 | 0 | 1 | 0 | 1 | 0 | 1 |
| B3 | 1 | 0 | 0 | 1 | 0 | 1 | 0 |
| B4 | 0 | 1 | 0 | 0 | 1 | 0 | 1 |
| B5 | 1 | 0 | 1 | 0 | 0 | 1 | 0 |
| B6 | 0 | 1 | 0 | 1 | 0 | 0 | 1 |
| B7 | 1 | 0 | 1 | 0 | 1 | 0 | 0 |
| B8 | 1 | 0 | 1 | 0 | 1 | 0 | 1 |
| B9 | 1 | 1 | 0 | 1 | 0 | 1 | 0 |
| B10 | 0 | 1 | 1 | 0 | 1 | 0 | 1 |
| B11 | 1 | 0 | 1 | 1 | 0 | 1 | 0 |
| B12 | 0 | 1 | 0 | 1 | 1 | 0 | 1 |
| B13 | 1 | 0 | 1 | 0 | 1 | 1 | 0 |
| B14 | 0 | 1 | 0 | 1 | 0 | 1 | 1 |

**Figure S5: Steady states of the odd-numbered toggle polygons with self-activation**. Frequency of most dominant states of the odd-numbered element circuits with self-activation added on each node as received from the RACIPE and Boolean simulations. **A.** 5cS. **B.** 7cS 0 and 1 respectively represents lower and higher concentration of corresponding component in the steady state solution of the corresponding state. N=3 independent RACIPE replicated were done; error bars denote standard deviation.

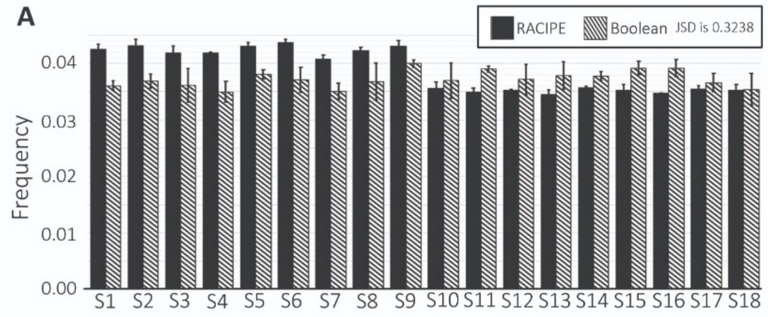


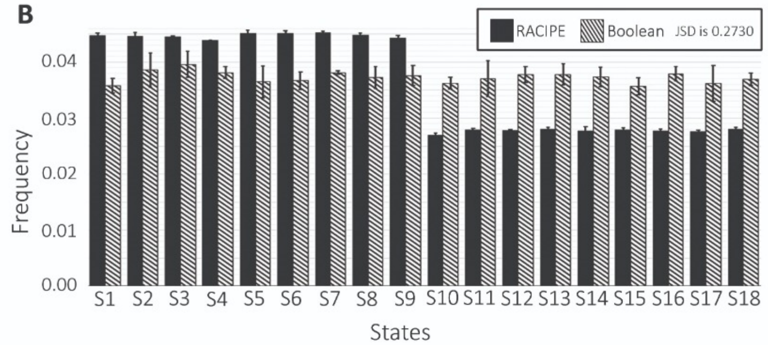


| State | A | B | C | D | E | F | G | H | I |
|---|---|---|---|---|---|---|---|---|---|
| S1 | 0 | 1 | 0 | 1 | 0 | 1 | 0 | 1 | 0 |
| S2 | 0 | 0 | 1 | 0 | 1 | 0 | 1 | 0 | 1 |
| S3 | 1 | 0 | 0 | 1 | 0 | 1 | 0 | 1 | 0 |
| S4 | 0 | 1 | 0 | 0 | 1 | 0 | 1 | 0 | 1 |
| S5 | 1 | 0 | 1 | 0 | 0 | 1 | 0 | 1 | 0 |
| S6 | 0 | 1 | 0 | 1 | 0 | 0 | 1 | 0 | 1 |
| S7 | 1 | 0 | 1 | 0 | 1 | 0 | 0 | 1 | 0 |
| S8 | 0 | 1 | 0 | 1 | 0 | 1 | 0 | 0 | 1 |
| S9 | 1 | 0 | 1 | 0 | 1 | 0 | 1 | 0 | 0 |
| S10 | 1 | 0 | 1 | 0 | 1 | 0 | 1 | 0 | 1 |
| S11 | 1 | 1 | 0 | 1 | 0 | 1 | 0 | 1 | 0 |
| S12 | 0 | 1 | 1 | 0 | 1 | 0 | 1 | 0 | 1 |
| S13 | 1 | 0 | 1 | 1 | 0 | 1 | 0 | 1 | 0 |
| S14 | 0 | 1 | 0 | 1 | 1 | 0 | 1 | 0 | 1 |
| S15 | 1 | 0 | 1 | 0 | 1 | 1 | 0 | 1 | 0 |
| S16 | 0 | 1 | 0 | 1 | 0 | 1 | 1 | 0 | 1 |
| S17 | 1 | 0 | 1 | 0 | 1 | 0 | 1 | 1 | 0 |
| S18 | 0 | 1 | 0 | 1 | 0 | 1 | 0 | 1 | 1 |

**Figure S6:** Frequency of most dominant states of the 9 element networks with (9cS) and without (9c) self-activation added on each node as received from the RACIPE and Boolean simulations. **A.** 9c **B.** 9cS. 0 and 1 respectively represents lower and higher concentration of the corresponding component in the steady state solution of the corresponding state. N=3 independent RACIPE replicates were done; error bars denote standard deviation.

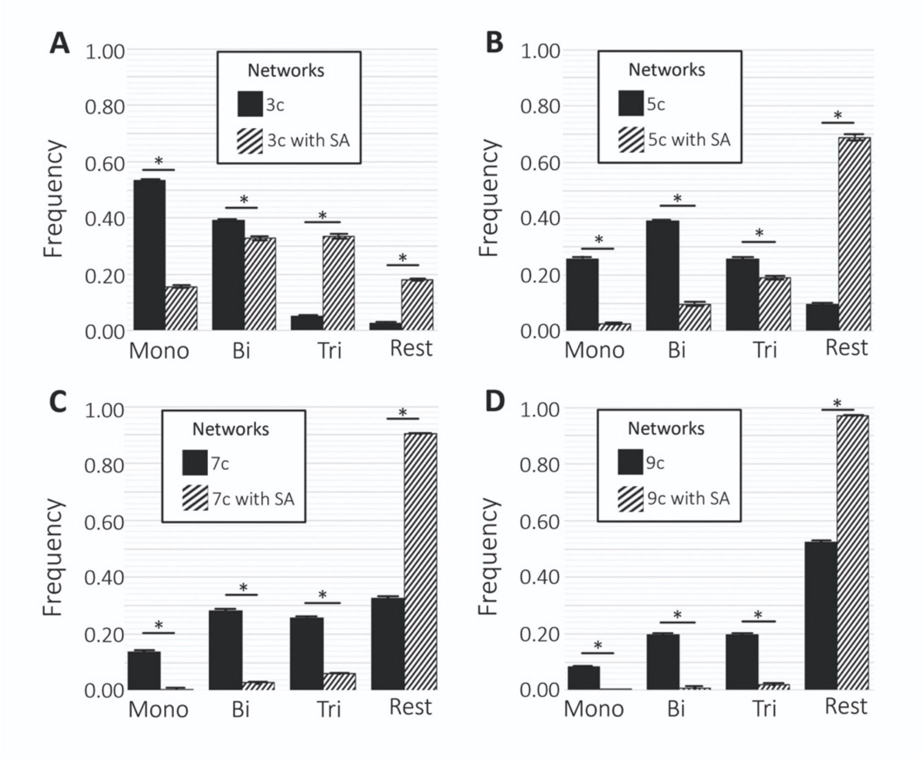


**Figure S7: Multistability in odd-numbered toggle polygons.** Comparison of frequencies of the monostable, bistable and tristable solutions form the RACIPE simulations of the **odd** element toggle polygon networks with and without added self-activation at each node. **A.** 3c and 3cS. **B.** 5c and 5cS. **C.** 7c and 7cS. **D.** 9c and 9cS. N=3 independent RACIPE replicated were done; error bars denote standard deviation. * denotes p<0.01 for a Student's t-test.

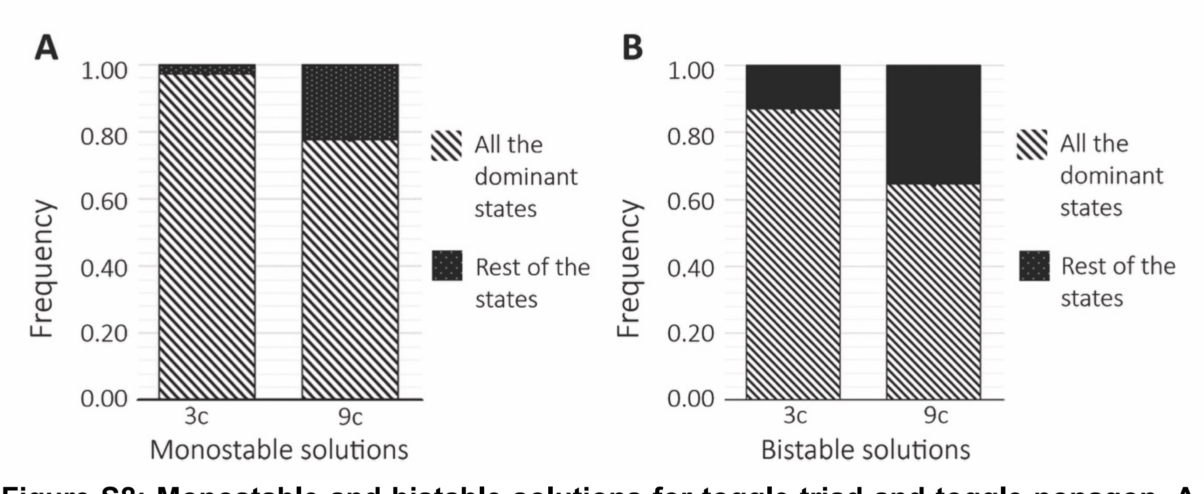


**Figure S8: Monostable and bistable solutions for toggle triad and toggle nonagon. A.** Frequency of all dominant monostable states combined with respect to the whole monostable solution space for the RACIPE simulations of 3c and 9c networks. **B.** Combined frequency of all the bistable states that are combinations of the most dominant monostable states with respect to the whole bistable state solution space for RACIPE simulations of 5c and 7c.N=3 independent RACIPE replicated were done; error bars denote standard deviation.

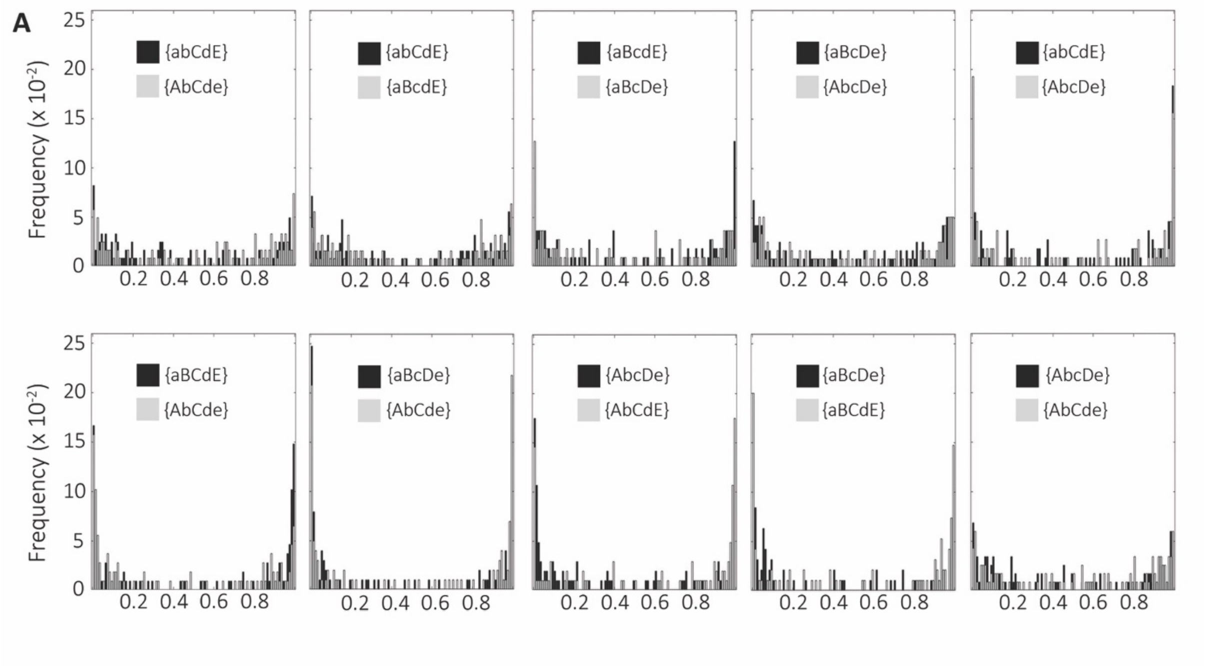


**Figure S9: Relative stability in toggle pentagon. A.** Relative frequency of corresponding monostable states in most dominant bistable solutions 5 element toggle polygon networks(5c) using the RACIPE generated parameters.

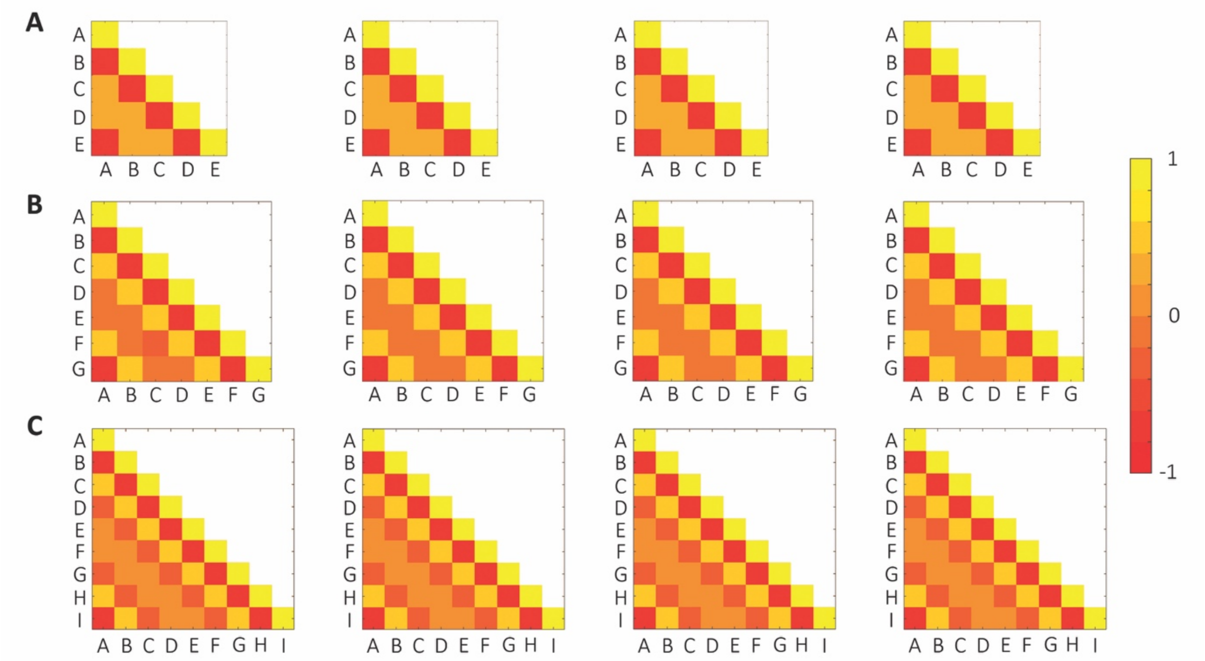


**Figure S10: Pairwise correlation matrices for odd-numbered toggle polygons.** (Left to right) Pearson correlation coefficients of monostable RACIPE solutions, Spearman correlation coefficients of monostable RACIPE solutions, Pearson correlation coefficients of all RACIPE solutions combined, and Spearman correlation coefficient of all RACIPE solutions combined on circuit topologies **A.** 5c, **B.** 7c, **C.** 9c. Each cell in this triangular matrix represents pairwise correlation. Colorbar shown to the left shows the value of correlation coefficients.

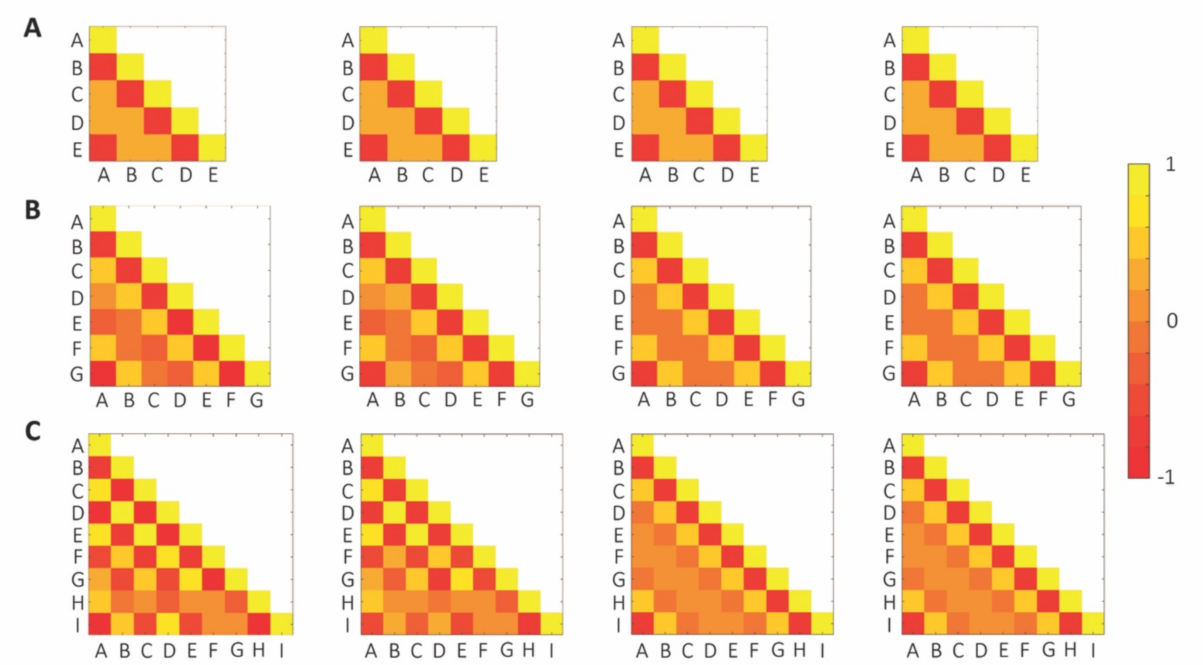


**Figure S11: Pairwise correlation matrices for even-numbered toggle polygons with self-activation.** Same as Fig S10 but for **A.** 5cS (toggle pentagon with self-activation), **B.** 7cS (toggle heptagon with self-activation), **C.** 9cS (toggle nonagon with self-activation).
